# Peroxisomes move by hitchhiking on early endosomes using the novel linker protein PxdA

**DOI:** 10.1101/034231

**Authors:** John Salogiannis, Martin J. Egan, Samara L. Reck-Peterson

**Affiliations:** Department of Cell Biology, Harvard Medical School, Boston, Massachusetts, 02115, USA; Department of Cellular and Molecular Medicine, University of California San Diego, La Jolla, CA 92093, USA; Section of Cell and Developmental Biology, Division of Biological Sciences, University of California San Diego, La Jolla, CA 92093, USA

## Abstract

Eukaryotic cells use microtubule-based intracellular transport for the delivery of many subcellular cargos, including organelles. The canonical view of organelle transport is that organelles directly recruit molecular motors via cargo-specific adaptors. In contrast to this view, we show here that peroxisomes move by hitchhiking on early endosomes, an organelle that directly recruits the transport machinery. Using the filamentous fungus *Aspergillus nidulans* we find that hitchhiking is mediated by a novel endosome-associated linker protein, PxdA. PxdA is required for normal distribution and long-range movement of peroxisomes, but not early endosomes or nuclei. Using simultaneous time-lapse imaging we find that early endosome-associated PxdA localizes to the leading edge of moving peroxisomes. We identify a coiled-coil region within PxdA that is necessary and sufficient for early endosome localization and peroxisome distribution and motility. These results present a new mechanism of microtubule-based organelle transport where peroxisomes hitchhike on early endosomes and identify PxdA as the novel linker protein required for this coupling.

## Introduction

Eukaryotic cells rely on the microtubule cytoskeleton to move intracellular components over long distances. Microtubules are dynamic polar structures, with “plus” ends usually located near the cell periphery and “minus” ends typically embedded in peri-nuclear microtubule organizing centers. Dynein motors move cargos towards the minus ends of microtubules, whereas most kinesin motors move in the opposite direction. Cytoplasmic dynein-1 (“dynein” here) and a relatively small number of kinesins are responsible for the transport of vesicles, organelles, proteins, and mRNAs (Cianfrocco et al., 2015; Vale, 2003).

In mammalian cells a variety of organelles have been shown to be dependent on dynein and kinesin for transport including endosomes, mitochondria, peroxisomes, Golgi, endoplasmic reticulum, autophagosomes, lysosomes, and nuclei (Harada et al., 1998; Kural et al., 2005; Maday and Holzbaur, 2012; Neuhaus et al., 2015; Roghi and Allan, 1999; Schrader et al., 2000; Tanaka et al., 1998). In some cases the adaptors that link the molecular motors to their cargos have been identified (Cianfrocco et al., 2015; Fu and Holzbaur, 2014; Kardon and Vale, 2009). For example, in the case of mitochondria, TRAK/Milton proteins recruit kinesin-1 and dynein (Glater et al., 2006; van Spronsen et al., 2013; Wang and Schwarz, 2009), and in the case of early endosomes Hook proteins have been shown to recruit dynein and kinesin-3 (Bielska et al., 2014; Zhang et al., 2014). This has led to the idea that each type of cargo uses distinct machinery to recruit molecular motors. Our goal is to use the model fungus *Aspergillus nidulans* to identify how distinct cargos engage the transport machinery.

*A. nidulans* and other filamentous fungi have proven to be excellent model systems for studying microtubule-based transport (Egan et al., 2012a). In these fungi, long-range microtubule-based transport is used to transport cellular cargos through highly polarized, multinucleate cells, called hyphae. Furthermore, rapid forward genetics are possible and genome engineering is fast and simple (Horio and Oakley, 2005; Nayak et al., 2006). In *A. nidulans*, microtubules are uniformly polarized from the hyphal tip to the tip-proximal nucleus, such that plus ends are located at hyphal tips and minus ends are embedded in the tip-proximal nuclear membrane (Egan et al., 2012b). As a result of this organization, defects in dynein-mediated cargo transport generally lead to accumulation of cargo at the hyphal tip, while defects in kinesin-3/UncA cargo transport can cause accumulation of cargo at the tip-proximal nucleus (see Fig. 1) (Egan et al., 2012a).

**Figure 1.**
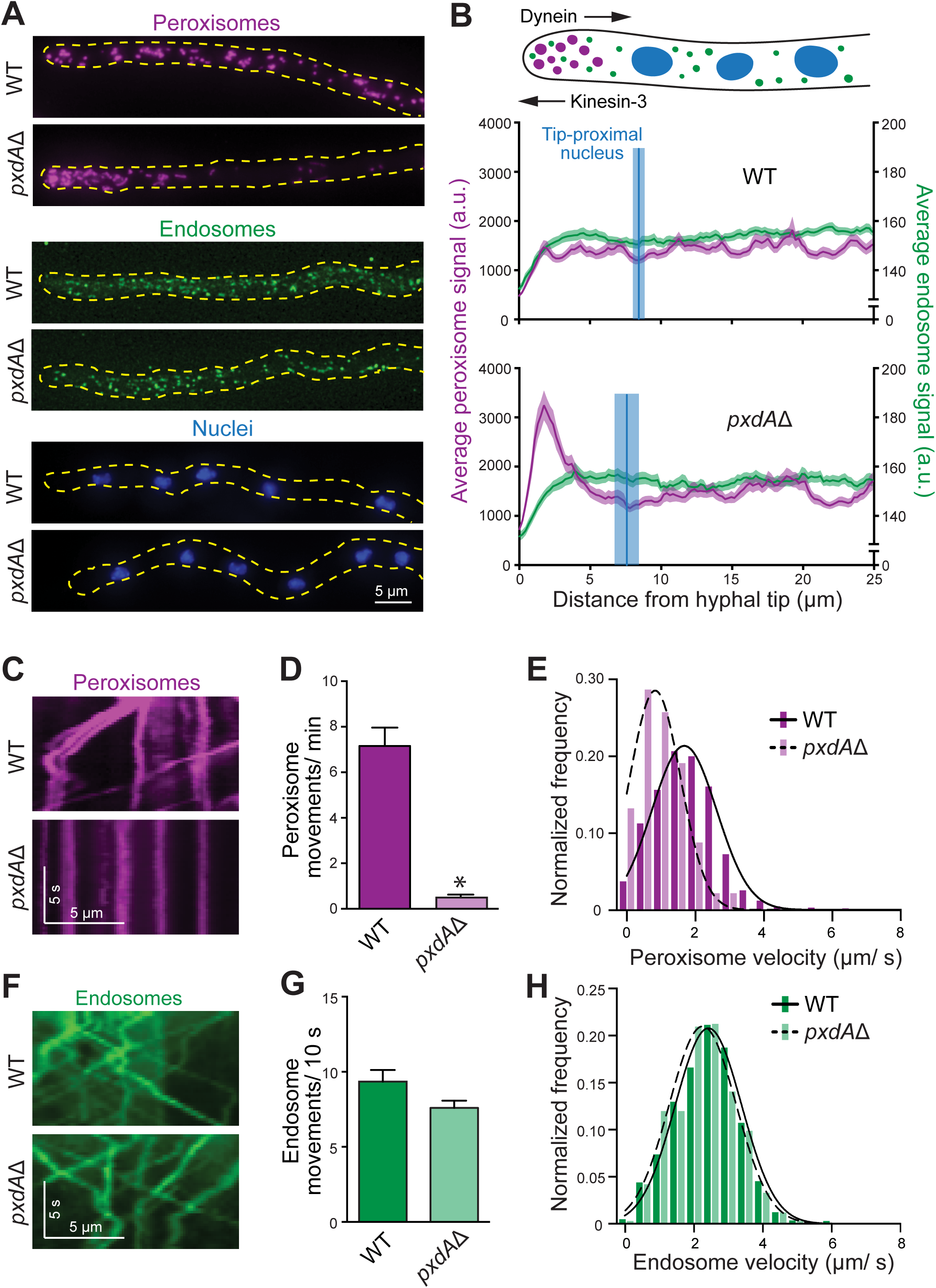
*pxdA* is required for the distribution and movement of peroxisomes, but not endosomes or nuclei. A) Representative micrographs of wild-type (WT) and *pxdA*Δ hyphae expressing fluorescent proteins that label peroxisomes (mCherry-peroxisome targeting signal 1 [PTS1]), early endosomes (GFP-RabA/5a) or nuclei (Histone H1 [HH1]-TagGFP). B) Cartoon of an *A. nidulans* hypha (top). Dynein moves away from the hyphal tip, while kinesin-3 moves in the opposite direction. Peroxisome (magenta) and early endosome (green) distribution in hyphae was quantified using line scans of fluorescence micrographs and displayed as the mean (solid lines) +/− SEM (shading) fluorescence intensity as a function of distance from the hyphal tip. Peroxisome distribution near the hyphal tip was significantly different between WT and *pxdA*Δ (p < 0.001, two-way ANOVA, Bonferroni post-hoc test significant between 0.65 μm and 3.25 μm from hyphal tip, n = 40 [WT], n = 44 [*pxdA*Δ]). Endosome distribution was not significantly different between WT and *pxdA*Δ (p = 0.99, two-way ANOVA, n = 51 [WT], n = 33 [*pxdA*Δ]). Nuclear distribution was quantified by measuring the distance from the distal nucleus to the hyphal tip. The mean (blue vertical lines) +/− SEM (blue shading) was 8.47 +/− 0.39 μm in WT and 7.66 +/− 0.42 μm in *pxdA*Δ hyphae. The means were not significantly different between WT and *pxdA*Δ (p = 0.1073, Mann Whitney test, n = 31 [WT], n = 31 [*pxdA*Δ]). C) Representative kymographs generated from time-lapse movies of mCherry-PTS1-labeled peroxisomes in WT and *pxdA*Δ hyphae. D) Bar graph of the flux of peroxisome movements in WT and *pxdA*Δ hyphae, calculated as the number of peroxisomes crossing a line drawn perpendicular and 10 μm away from the hyphal tip during a 1-min time-lapse movie. Peroxisome movements were 7.17 +/− 0.79 (SEM)/ min in WT and 0.52 +/− 0.10/ min in *pxdA*Δ hyphae (*p < 0.0001, Mann Whitney test, n = 44 for both genotypes). E) Histogram of instantaneous velocities (binned in 0.5 μm increments) of moving peroxisomes (defined as greater than 2 μm displacement during a 1-min time-lapse video) in WT versus *pxdA*Δ strains. Mean velocities were 1.75 +/− 0.99 (SD) μm/ s in WT and 0.97 +/− 0.68 μm/ s in *pxdA*Δ hyphae (p < 0.001, Kolmogorov-Smirnov test, n = 1195 [WT], n = 136 [*pxdA*Δ]). F) Kymographs generated from time-lapse movies of GFP-RabA/5a -labeled endosomes in WT and *pxdA*Δ hyphae. G) Bar graph of the flux of endosome movements during a 10 s time-lapse movie calculated as in D). Endosome movements were 9.34 +/− 0.78 (SEM)/ min in wild-type and 7.60 +/− 0.48/ min in *pxdA*Δ hyphae (p = 0.0920, Mann Whitney test, n = 29 [WT], n = 47 [*pxdA*Δ]). H) Histogram of instantaneous velocities of endosomes in WT versus *pxdA*Δ strains. Mean velocities are 2.40 +/− 0.99 (SD) μm/ s in WT and 2.24 +/− 0.93 μm/ s in *pxdA*Δ hyphae (p = 0.0274, Kolmogorov-Smirnov test, n = 1602 [WT], n = 332 [*pxdA*Δ]).

We took advantage of these clear organelle distribution phenotypes to conduct a forward genetic mutagenesis screen in *A. nidulans* to identify novel genes required for the microtubule-based movement of dynein and kinesin cargos (Tan et al., 2014). For our screen we focused on three well-characterized cargos of dynein and kinesin: early endosomes, peroxisomes, and nuclei (Abenza et al., 2009; Egan et al., 2012b; Wedlich-Soldner et al., 2002; Xiang et al., 1994). We identified novel alleles of known components of the transport machinery as well as many genes not previously connected to transport (Tan et al., 2014). One expectation was that some of these novel genes would encode organelle-specific adaptors for dynein or kinesin.

In the present study we focused on hits from our screen that resulted in deficits in the specific distribution and movement of peroxisomes. Microtubule-based movement of peroxisomes is likely critical to distribute these multi-functional organelles evenly throughout the cell and to guard against toxic reactive oxygen species and long-chain fatty acids (Liu et al., 2008; Neuhaus et al., 2015). We identified a novel coiled-coil-containing protein, PxdA, which is required for the distribution and long-range movement of peroxisomes. We show that PxdA colocalizes with early endosomes, and that its endosomal localization is required for peroxisome movement. We find that both PxdA and early endosomes co-migrate with moving peroxisomes. Recently, it was reported that peroxisomes, as well as other cargo, could “hitchhike” on early endosomes in the plant pathogenic fungus *Ustilago maydis* to achieve long-range movement (Baumann et al., 2012; Guimaraes et al., 2015; Higuchi et al., 2014), but the molecules mediating this behavior were not known. Our data demonstrate that PxdA is a novel linker protein mediating peroxisomal hitchhiking on early endosomes.

## Results and Discussion

### PxdA is a novel protein required for peroxisome distribution and motility

In our original screen 19 mutants were categorized as only affecting peroxisome distribution, making them excellent candidates for cargo-specific regulators (Tan et al., 2014). Whole-genome sequencing of two mutants (RPA604 and RPA628) with a strong peroxisome distribution phenotype identified independent mutations in the uncharacterized gene AN1156; both mutations were predicted to result in premature stop codons. AN1156 mutants displayed hyphal tip accumulation of peroxisomes, but normal distribution of RabA/5a-positive early endosomes (“endosomes” from here on) and nuclei (Fig. S1 A). Based on this phenotype we named AN1156 *pxdA* (***p***ero***x***isome ***d***istribution mutant ***A***). To confirm that PxdA is expressed we performed immunoblots of lysates from a HA-tagged *pxdA* strain and detected a ~250 kDa band (Fig. S1 B), which corresponds well to the larger of two predicted open reading frames for AN1156.

To verify that *pxdA* plays a role in regulating peroxisome distribution we deleted the endogenous gene in haploid strains containing fluorescently labeled endosomes, peroxisomes, or nuclei. Consistent with the phenotype of RPA604 and RPA628, peroxisomes accumulated in *pxdA*Δ hyphal tips, but endosomes and nuclei were distributed normally (Fig. 1 A). To quantify peroxisome and endosome distribution along hyphae, we performed line scans of fluorescence micrographs and plotted the average fluorescence intensity as a function of distance from the hyphal tip. The average peroxisome signal increased significantly in *pxdA*Δ hyphal tips, but no differences were observed in the average endosome signal or the average position of the tip-proximal nucleus (Fig. 1 B). As an independent measurement of peroxisome distribution, we found that peroxisomes labeled with PexK (a homolog of *S. cerevisiae* Pex11) (Hynes et al., 2008) also accumulated in the hyphal tips of *pxdA*Δ strains (Fig. S1, C and D). We conclude that PxdA specifically regulates the distribution of peroxisomes, but not endosomes or nuclei.

Peroxisome distribution and motility in *A. nidulans* requires the microtubule-based motors cytoplasmic dynein/NudA and kinesin-3a/UncA (Egan et al., 2012a; Egan et al., 2012b). To determine whether the loss of *pxdA* affects peroxisome motility as well as distribution we imaged peroxisomes in wild-type and *pxdA*Δ strains. In wild-type hyphae peroxisomes exhibit both long-range runs and oscillatory movements (Video 1). In wild-type hyphae, 12% (n = 486) of peroxisomes exhibit long-range movements (distances greater than 3 μm), consistent with previous studies in mammalian cells where only 10–15% of peroxisomes are motile (Bharti et al., 2011; Rapp et al., 1996). These long-range movements are reduced by greater than 90% in *pxdA*Δ hyphae (Fig. 1, C and D; Video 2). Furthermore, the velocity and run length of the small pool of peroxisomes that move in *pxdA*Δ hyphae are also reduced compared to wild-type (Fig. 1 E and Fig. S1 E). In contrast, the flux and velocity of endosomes are similar in wild-type and *pxdA*Δ strains (Fig. 1, F – H; Video 3 and 4). These findings are consistent with a specific role for PxdA in the long-range movements of peroxisomes.

Taken together, our findings thus far demonstrate that PxdA is required for the proper distribution and long-range movement of peroxisomes, but is not required for the distribution or motility of endosomes or nuclei. Our data also suggest that PxdA is not a core component of the transport machinery, since deletions of this type would be predicted to result in defects in the transport or distribution of endosomes and nuclei as well. Consistent with this, we find that the localization of dynein, characteristically seen as “comets” near the microtubule-plus-end (Xiang et al., 1995), is intact in *pxdA*Δ hyphae (Fig. S1 F).

### Peroxisomal hitchhiking on early endosomes is mediated by PxdA

Since PxdA is essential for peroxisome movement, we hypothesized that it might colocalize with moving peroxisomes. To visualize PxdA, we tagged the endogenous copy at its C-terminus with either TagGFP (“GFP” from here on) or the red fluorescent protein, mKate. C-terminal tagging of PxdA did not disrupt its function as line-scan analysis revealed that PxdA-GFP hyphae have normal peroxisome distribution (Fig. S2 A). In time-lapse videos, PxdA exhibits rapid bi-directional movement along the entire length of hyphae (Video 5). Kymograph analysis reveals that the velocity of PxdA is consistent with that of microtubule-based movement (Fig. 2, A and B). To evaluate the role of microtubules directly we treated the PxdA-GFP strain with benomyl, a drug that inhibits microtubule polymerization and dynamic instability. The flux of bidirectional PxdA movements is largely abolished in benomyl-versus DMSO-treated conditions (Fig. 2 C). In contrast, we found no defects in PxdA flux in the presence of the actin polymerization inhibitor, latrunculin-A (Fig. 2 C).

**Figure 2.**
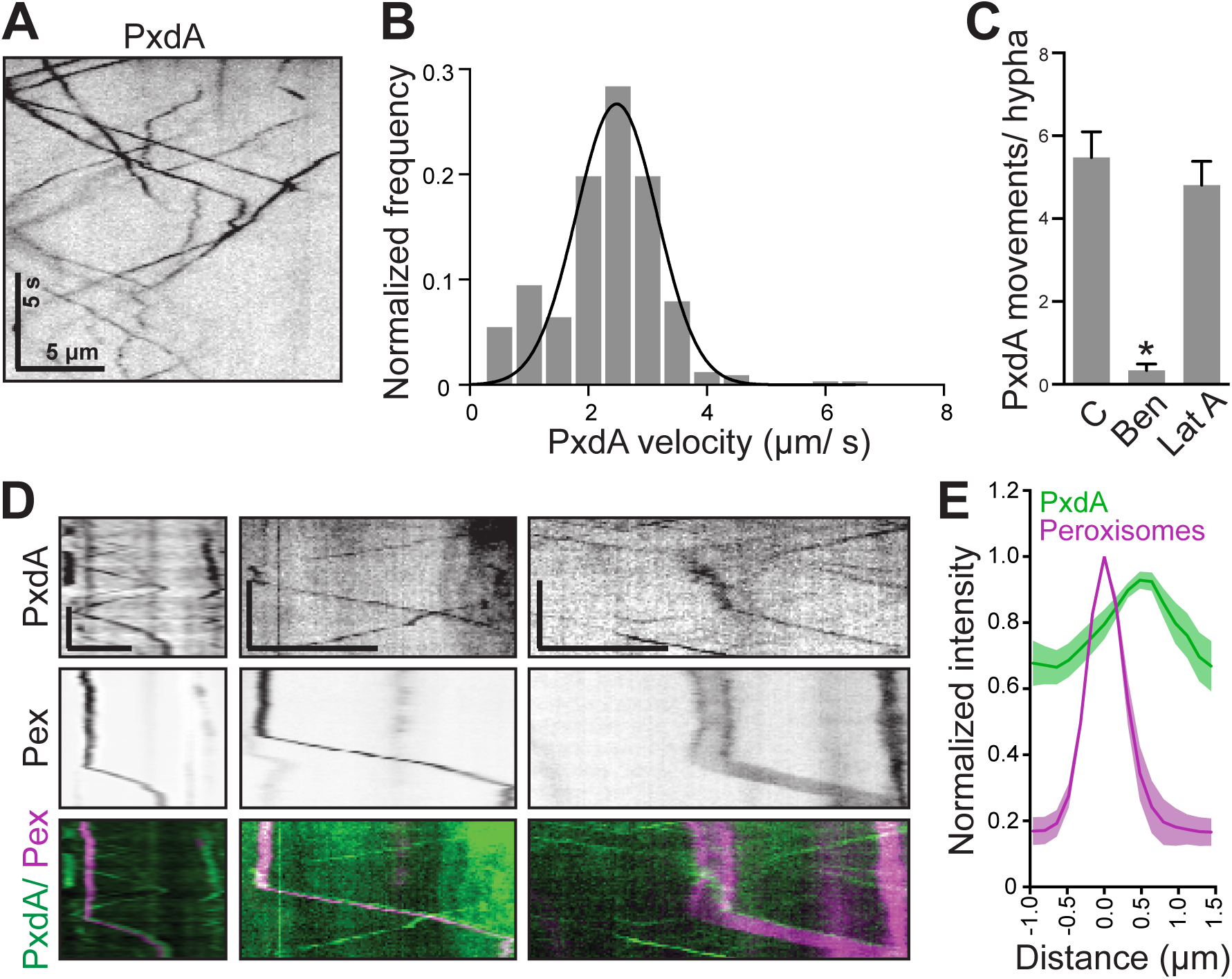
Microtubule-based movement of PxdA colocalizes with moving peroxisomes. A) Representative kymograph generated from a time-lapse movie of PxdA-GFP. B) Histogram of velocities calculated from PxdA-GFP and PxdA-mKate kymographs. Mean velocity is 2.32 +/− 0.88 (SD) (n = 328). C) Bar graph of the number of PxdA-mKate movements calculated as the number of puncta crossing a line drawn perpendicular to the hyphal long axis during a 10 s movie. PxdA puncta moved 5.48 +/− 0.62 (SEM)/ hyphae in the DMSO control (C, n = 21), 0.33 +/− 0.16/ hyphae in the presence of benomyl (Ben, n = 15), and 4.81 +/− 0.57/ hyphae in the presence of latrunculin A (LatA, n = 16). Benomyl, but not latrunculinA treatment was significantly different than the control (one-way ANOVA, Bonferonni post-hoc test, *p < 0.001). D) Representative kymographs generated from simultaneous, dual-color, time-lapse movies of PTS1-mCherry labeled peroxisomes (pex) and PxdA-GFP. 65.6% (n = 29) of moving peroxisomes overlapped with PxdA. Scale bars are 5 μm (x axis) and 5 s (y axis). Left panel corresponds to Video 5. E) Quantification of intensity line scans of peroxisomes (magenta) and PxdA (green) during comigrating runs. Normalized peak intensity of PxdA was 0.56 +/− 0.08 (SEM) μm from the normalized peak intensity of peroxisomes (n = 11).

To determine if PxdA colocalizes with peroxisomes we next visualized PxdA and peroxisomes simultaneously using dual-color time-lapse imaging. Kymograph analysis (Fig. 2 D) revealed that 66% (n = 29 moving peroxisomes) of moving peroxisomes co-migrate with PxdA. PxdA is rarely associated with stationary peroxisomes. Closer inspection of the time-lapse movies and kymographs revealed that PxdA localizes at the leading edge of moving peroxisomes, with the PxdA signal leading by 560 +/− 80 nm (SEM) (Fig. 2 E; Video 6). We also occasionally observed instances where a moving PxdA puncta paused at a nonmotile peroxisome, an event that was followed by the movement of co-migrating PxdA and the peroxisome (Fig. 2 D middle and right panels).

Because we observed many more moving PxdA puncta than moving peroxisomes and because PxdA motility was reminiscent of endosome motility (Abenza et al., 2009; Egan et al., 2015; Egan et al., 2012b; Zhang et al., 2010), we next asked whether PxdA also colocalizes with endosomes. To test this, we analyzed kymographs from simultaneous time-lapse movies of labeled PxdA and endosomes (Video 7). Ninety percent of PxdA puncta colocalize with endosomes, but only 55% of endosomes colocalize with PxdA puncta (Fig. 3, A and B), demonstrating that PxdA is found on a subset of endosomes.

**Figure 3.**
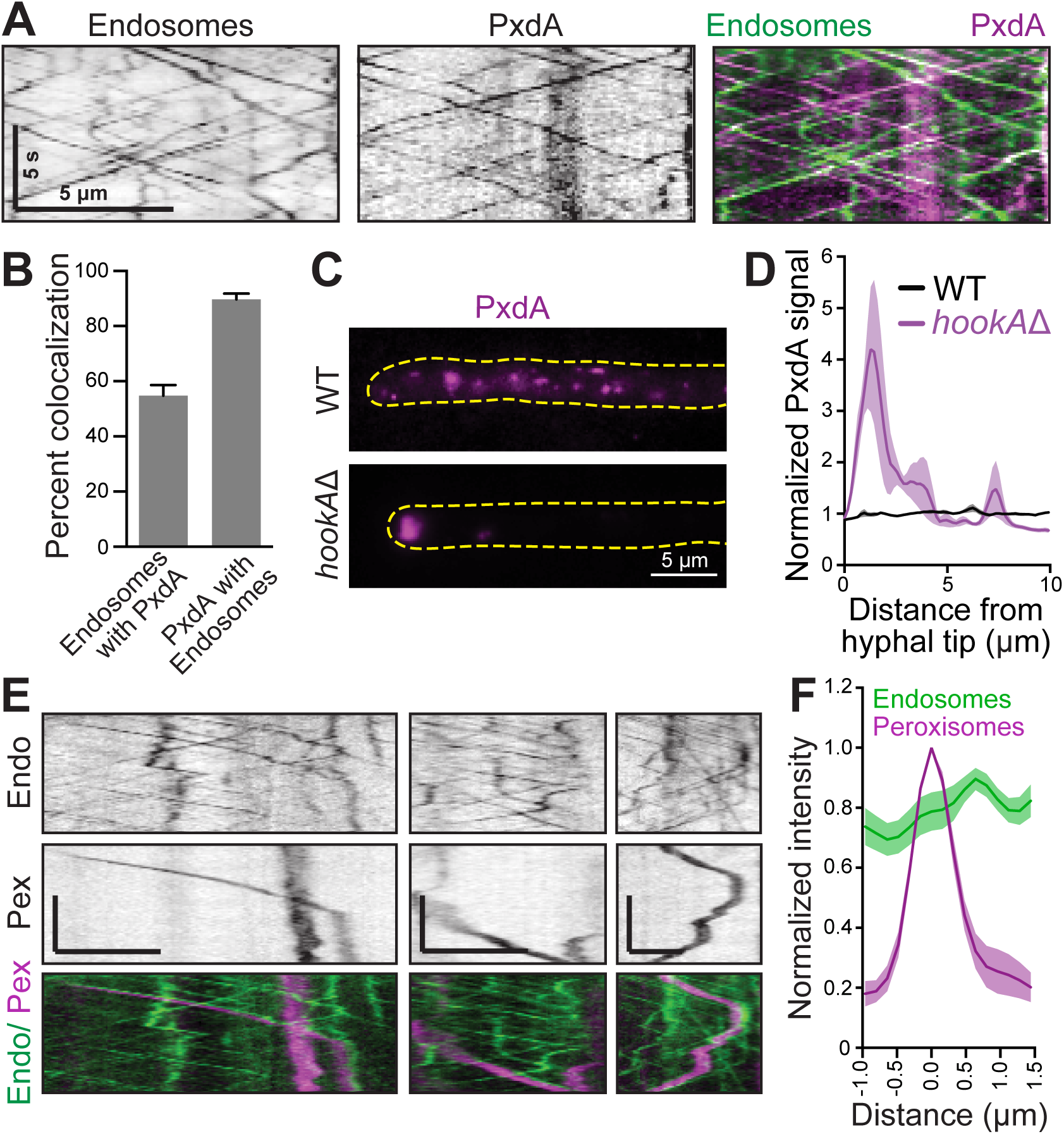
Early endosomes colocalize with PxdA and moving peroxisomes. A) Representative kymographs generated from simultaneous, dual-color time-lapse movies of PxdA-mKate and GFP-RabA/5a-labeled endosomes. (also see Video 7). B) Bar graph quantifying the colocalization of PxdA-mKate and endosomes. Endosomes colocalized with 54.8 +/− 3.8% (SEM) PxdA puncta (n = 20 kymographs from 10 cells). PxdA puncta colocalized with 89.7 +/− 2.0% endosomes (n = 20 kymographs from 10 cells). C) Representative micrographs of PxdA-mKate distribution in wild-type versus *hookA*Δ hyphae. D) Normalized line scans of WT and *hookA*Δ hyphae. (p < 0.0001, two-way ANOVA, Bonferonni post-hoc test significant between 0.64 and 1.92 μm, n=5 per genotype) E) Representative kymographs generated from simultaneous, time-lapse movies of PTS1-mCherry-labeled peroxisomes (pex) and GFP-RabA/5a-labeled endosomes (endo). 71.4% (n = 28) of moving peroxisomes overlapped with early endosomes. Scale bars are 5 μm (x axis) and 5 s (y axis). Left panel corresponds to Video 8. F) Quantification of intensity line scans of peroxisomes (magenta) and endosomes (green) during comigrating runs. Normalized peak intensity of endosomes was 0.61 +/− 0.17 (SEM) μm from the normalized peak intensity of peroxisomes (n = 11).

We next sought to determine if perturbing endosome localization also perturbed PxdA localization. HookA has recently been identified as a cargo adaptor between early endosomes and cytoplasmic dynein and kinesin-3 in filamentous fungi (Bielska et al., 2014; Zhang et al., 2014), and its vertebrate homolog (Hook3) is an activator of dynein/dynactin (McKenney et al., 2014). In the absence of HookA early endosomes accumulate near the hyphal tip (Bielska et al., 2014; Zhang et al., 2014). We reasoned that if most PxdA protein is associated with early endosomes, PxdA would accumulate at hyphal tips in *hookA*Δ strains. Indeed, we find that PxdA accumulates at the hyphal tip in *hookA*Δ compared to wild-type strains (Fig. 3, C and D), suggesting that PxdA is present on endosomes that require HookA for motility.

The association between PxdA and endosomes is somewhat surprising given that PxdA does not affect the motility or distribution of endosomes (Fig. 1). Since PxdA colocalizes with a subset of early endosomes and moving peroxisomes, we next asked if endosomes colocalized with moving peroxisomes. Kymograph analysis (Fig. 3 E) of time-lapse movies reveals that 71% (n = 28) of moving peroxisomes colocalize with endosomes, correlating well with the percentage of PxdA that colocalizes with moving peroxisomes (66%). In addition, similar to PxdA, endosomes colocalized at the leading edge of moving peroxisomes (Fig. 3 F; Video 8). The endosome signal was leading the peroxisome signal by 610 +/− 170 nm (mean +/− SEM), a number comparable to what we observed for peroxisomes and PxdA (Fig. 2 E).

Taken together, these results suggest that peroxisomes move by “hitchhiking” on early endosomes and that this process is mediated by PxdA. The concept that cellular components can hitchhike on endosomes has been observed in *U. maydis* for Rrm4-containing ribonucleoprotein complexes (Baumann et al., 2012) and polysomes (Higuchi et al., 2014). In a parallel study to ours, also in *U. maydis*, peroxisomes, endoplasmic reticulum and lipid droplets were observed to hitchhike on early endosomes (Guimaraes et al., 2015).

### Endosomal localization of PxdA is required for peroxisome hitchhiking

We hypothesized that PxdA tethers endosomes to peroxisomes to promote hitchhiking. If this were the case, we might be able to separate PxdA’s endosomal localization from its function in peroxisome motility. To test this, we constructed a series of PxdA mutants (Fig. 4 A).

**Figure 4.**
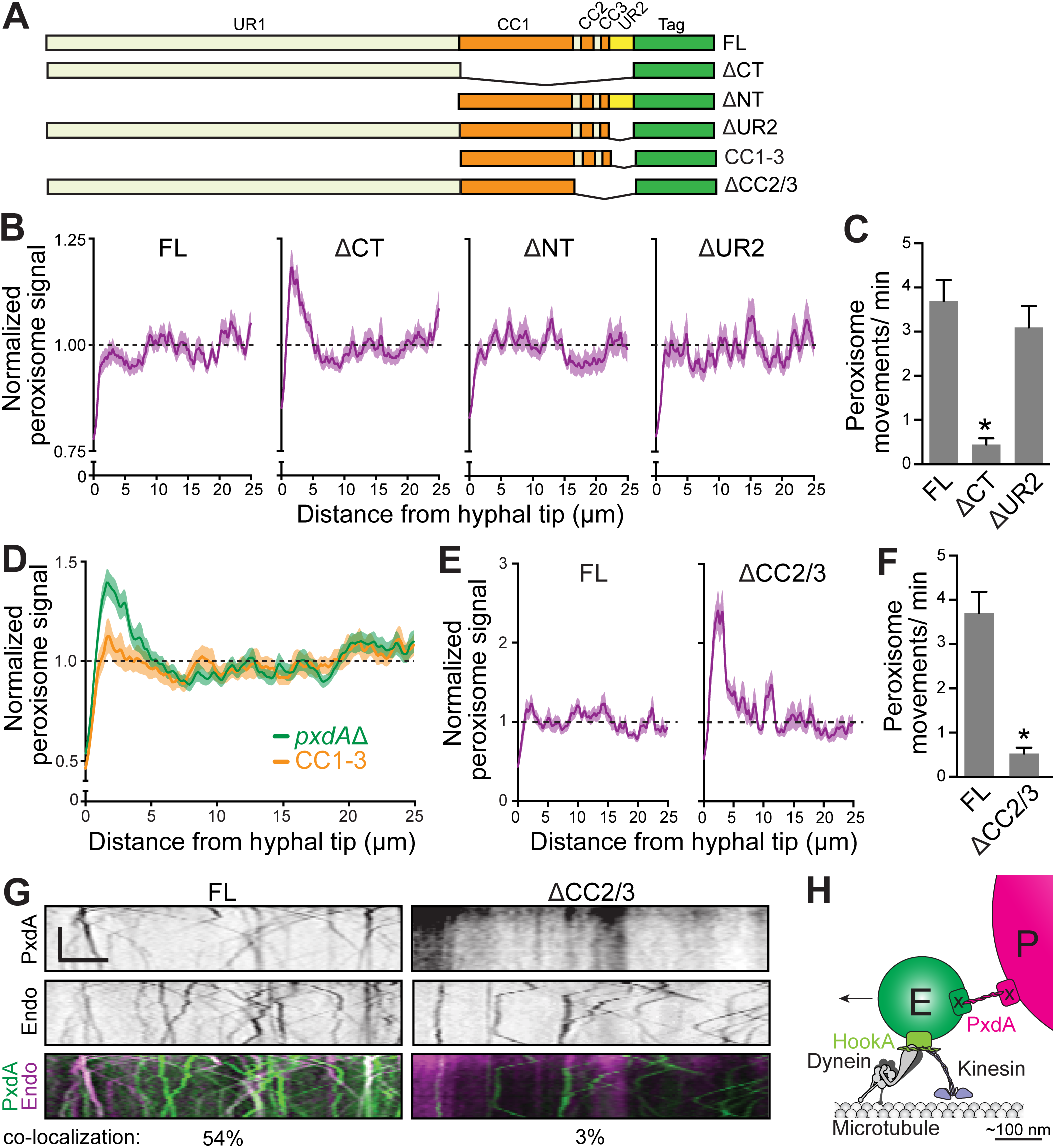
The coiled-coil region of PxdA is necessary and sufficient for peroxisome motility and endosomal hitchhiking. A) Schematic of PxdA protein constructs used in this analysis relative to full-length (FL) PxdA. Uncharacterized regions 1 (UR1, pale yellow) and 2 (UR2, yellow) are on the N- and C-terminus, respectively. Predicted regions of coiled-coil (CC1, 2, and 3) are indicated in orange. All strains contain a C-terminal fluorescent protein tag (either mKate or TagGFP) for visualization. B) Normalized line scans of peroxisome (PTS1-mCherry) distribution from the hyphal tip for FL, C-terminal deletion (ΔCT), N-terminal deletion (ΔNT) and ΔUR2 PxdA strains (p < 0.0001, two-way ANOVA, Bonferroni post-hoc test significant between 0.51 μm and 3.71 μm for FL vs ΔCT, n = 51–54 per genotype). C) Bar graph of the flux of peroxisome movements in FL, ΔCT, and ΔUR2 hyphae. Peroxisome movements were 3.69 +/− 0.47 (SEM)/ min for FL, 0.44 +/− 0.14/ min for ΔCT, 3.10 +/− 0.50/ min for ΔUR2. (p < 0.0001, one-way ANOVA, Bonferroni post-hoc test significant (*) for FL vs. ΔCT; FL vs. ΔUR2 was not significantly different, n = 29 − 35 per genotype) D) Normalized line scans of peroxisome distribution along hyphae expressing either FL or CC1–3 PxdA (p < 0.001, two-way ANOVA, Bonferroni post-hoc test significant between 1.19 μm and 3.03 μm from hyphal tip; n = 78 [FL], n = 68 [CC1–3]). E) Normalized line scans of peroxisome distribution along hyphae expressing either FL or ΔCC2/3 PxdA. (p < 0.0001, two-way ANOVA, Bonferroni post-hoc test significant between 0.87 and 4.11 μm from hyphal tip; n = 28 [FL], n = 32 [ΔCC2/3]. F) Bar graph of the flux of peroxisome movements in FL and ΔCC2/3 hyphae. Peroxisome movements were 3.69 +/− 0.47 (SEM)/ min for FL (this is the same FL data depicted in C) and 0.51 +/− 0.13/ min for ΔCC2/3 (p < 0.0001, one-way ANOVA, Bonferroni post-hoc test significant (*) for FL vs. ΔCC2/3, n = 29 − 35 per genotype) G) Representative kymographs generated from simultaneous time-lapse movies of GFP-RabA/5a-labeled endosomes with FL or ΔCC2/3 PxdA. Colocalization represents the number of endosomes that colocalize with PxdA. FL is 54.8% +/− 3.8% (from Fig. 3B) and ΔCC2/3 3.2% +/− 1.3% (n = 10 kymographs). H) Model for peroxisome (P) hitchhiking on early endosomes (E) mediated by PxdA. HookA recruits the transport machinery to early endosomes. PxdA is required to tether endosomes to peroxisomes and may have distinct binding partners (x) on each organelle. Arrow depicts the direction of movement.

First, we sought to determine which domains of PxdA are necessary and sufficient for peroxisome motility. PxdA encodes a 2236 amino acid protein containing a ~600 amino acid stretch of predicted tandem coiled-coil domains (CC1, 2 and 3), flanked by a large N-terminal uncharacterized region (UR1) and a much smaller C-terminal uncharacterized region (UR2) (Fig. 4 A). Initially, we truncated the entire N-terminal UR1 (ΔNT) or the entire C-terminal region including CC1-3 and UR2 (ΔCT), or the UR2 region alone (ΔUR2). We performed peroxisome line-scans and found that peroxisomes accumulated at the hyphal tip of ΔCT strains, but not full-length, ΔUR2 or ΔNT strains (Fig. 4 B). PxdA-ΔCT and ΔUR2 strains also had a significant reduction in the flux of peroxisome movements compared to the full-length strain (Fig. 4 C).

This analysis suggested that only the tandem coiled-coil domain (CC1-3) of PxdA is required for peroxisome distribution and motility. To test this directly, we expressed PxdA CC1-3 and quantified peroxisome distribution along hyphae. CC1-3 alone is sufficient to rescue the peroxisome accumulation observed in PxdAΔ hyphae (Fig. 4 D). PxdA CC1-3 localizes to moving puncta that show similar velocity and flux to strains expressing full-length PxdA (Fig. S2, B – D). These experiments show that CC1-3 is necessary and sufficient for PxdA’s function in peroxisome distribution.

We next sought to identify the region of PxdA required for its localization to endosomes. We constructed a PxdA-mKate strain lacking CC2 and CC3 (ΔCC2/3; Fig. 4 A). Like PxdA ΔCT strains, PxdA ΔCC2/3 strains also displayed an accumulation of peroxisomes at the hyphal tip (Fig. 4 E) and a severe drop in peroxisome flux (Fig. 4 F). In addition, PxdA ΔCC2/3-mKate had a diffuse signal in hyphae that was different from the endosome localization observed for the full-length protein and only 3% of PxdA ΔCC2/3-mKate colocalized with endosomes (Fig. 4 G). This data suggests that CC2/3 is critical for the endosome localization of PxdA, and that endosomally-linked PxdA is required for the proper distribution and movement of peroxisomes.

Overall, our study reveals that peroxisomes require PxdA to hitchhike on early endosomes for long-range movement. In addition to their role in endocytosis, the abundance and rapid bidirectional movement of early endosomes in fungal hyphae make them a prime platform for distributing a wide-range of cargo (Gohre et al., 2012). In our model, HookA recruits the transport machinery to early endosomes (Bielska et al., 2014; Zhang et al., 2014) and PxdA tethers peroxisomes to endosomes (Fig. 4 H). The observation that both PxdA and endosomes lead peroxisomes by ~600 nm during movement (Fig. 2 E and Fig. 3 E) is consistent with the approximate diameter of peroxisomes (~1 um, Egan et al., 2012b) and endosomes (~150 nm) (Griffith et al., 2011; Murk et al., 2003). The size of PxdA’s tandem coiled-coil (CC1-3), which is predicted to be ~90 nm could easily bridge the distance between the peroxisome and endosome. Exciting future directions include determining if hitchhiking is broadly conserved outside of fungi, identifying the linkers required for other cargos to hitchhike on early endosomes, and determining the mechanisms that initiate and terminate hitchhiking.

## Materials and Methods

### Fungal growth conditions

*A. nidulans* strains were grown in yeast extract and glucose (YG) medium (Szewczyk et al., 2006) or 1% glucose minimal medium (Nayak et al., 2006), supplemented with 1 mg/ ml uracil, 2.4 mg/ ml uridine, 2.5 μg/ ml riboflavin, 1 μg/ml para-aminobenzoic acid, and 0.5 μg/ ml pyridoxine when required. Glufosinate was used at a final concentration of 25 μl/ mL as previously described (Nayak et al., 2006).

For imaging of germlings, spores were resuspended in 0.5 mL 0.01% Tween-80 solution. The spore suspension was diluted at 1:1,000 in liquid minimal medium containing appropriate auxotrophic supplements. The spore and media mix (400 μL) was added to an eight-chambered Nunc Lab-Tek II coverglass (ThermoFisher) and incubated at 30°C for 16–20 hours before imaging. For imaging of mature hyphae, spores were inoculated on minimal medium plates containing the appropriate auxotrophic supplements and incubated at 37°C for 12–16 hours. Colonies were excised from agar plates and inverted on Lab-Tek plates for imaging. For benomyl and latrunculinA experiments, minimal media containing the drugs (2.4 μ/ mL for benomyl [Sigma]; 12 μM for latrunculinA [Life Technologies]) or DMSO control (1.2%) was added fresh to Lab-Tek chambers containing germlings and were imaged between 45 mins to one hour later. For Western blots, cells were grown in YG medium for 16–20 hours, strained in miracloth (Millipore) and flash frozen in liquid nitrogen. For lysis, cells were ground in liquid nitrogen and boiled in 9M urea denaturing buffer (125 mM Tris-HCl pH 6.8, 9M Urea, 1mM EDTA pH7.0, 4% SDS, 10mM DTT, 10% beta-ME, 4% glycerol) prior to running on SDS-PAGE.

### Strain Construction

Strains of *A. nidulans* used in this study are listed in Table S1. All strains were confirmed by a combination of PCR and sequencing from genomic DNA isolated as previously described (Lee and Taylor, 1990). Strains were created by homologous recombination to replace the endogenous gene in strains lacking *ku70 (Nayak et al., 2006)* with *Afribo (Aspergillus fumigatus Ribo), AfpyrG (Aspergillus fumigatus pyrG), Afpyro (Aspergillus fumigatus pyro)*, or *bar* (Straubinger, 1992) as selectable markers. All linearized transforming DNA for deletion strains including *pxdA*Δ and *hookA*Δ were constructed by fusion PCR (Szewczyk et al., 2006). All PxdA DNA constructs contained a C-terminal codon-optimized fluorescent protein tag, either TagGFP2 (Subach et al., 2008) or mKate2 (Shcherbo et al., 2009) followed by PxdA’s native 3’ UTR and were inserted into the Blue Heron Biotechnology pUC vector at 5’ EcoRI and 3’ HindIII restriction sites using isothermal assembly (Gibson et al., 2009). Plasmid constructs were confirmed by sequencing. Linearized transforming DNA from plasmid constructs was created by PCR with flanking primers located at the 5’ and 3’ ends of homologous arms.

The amino acid positions for each PxdA construct in Figure 4 were as follows: FL (aa 1-2236), ΔNT (aa 1466-2236), ΔCT (aa 1 - 1465), ΔUR2 (aa 1 – 2116), and ΔCC2/3 (aa 1 – 1946). For the CC1-3 rescue experiment (Fig. 4 D and Fig. S1, B - D), the CC construct (aa 1466 – 2116) was inserted into a *pxdA*Δ strain at the endogenous locus.

### Whole-genome sequencing

Methods were described in detail in our previous study (Tan et al., 2014).

### Fluorescence microscopy

All images were collected at 22°C. For acquisition of the distribution and motility of peroxisomes (PTS1-mCherry and PexK-GFP) and for the distribution of endosomes (GFP-RabA/5a) and dynein (NudA-3xGFP), images were collected using a Plan Apo 60x/1.42 (Olympus) oil immersion objective on an epifluorescence Deltavision Core microscope (GE Healthcare). GFP was excited with a 488nm laser line (50mW) and collected with a 525/50 FITC emission filter. mCherry fluorescence was excited with the 561nm laser line (50mW) and collected with a 594/45 emission filter. Images were acquired with a PCO edge sCMOS camera (Kelkeim, Germany) and controlled with softWoRx software (GE Healthcare). For distribution images, 20 optical z-sections were collected with a step size of 0.5 μms. Hyphae (used for outlining cells) were imaged using brightfield.

Simultaneous multicolor time-lapse images were collected using a Plan Apo TIRF 60x/1.49 oil immersion objective on the Deltavision OMX Blaze V4 system (GE Healthcare). GFP and mKate2/mCherry were excited simultaneously with 488nm and 568nm diode laser lines, respectively. A BGR polychroic mirror was used to split emission light from fluorophores to different PCO Edge sCMOS cameras. Emission filter in front of both cameras was used to select appropriate wavelengths (528/48 and 609/37 for GFP and mKate/mCherry, respectively). Images were aligned with OMX image registration using softWoRx software and some images were deconvolved for display using the enhanced ratio method.

For endosome and PxdA motility assays, time-lapse images were collected using a Plan Apo TIRF 100x/1.49 oil immersion objective on an epifluorescence Nikon Ti-E motorized inverted scope with the Perfect focus system (Nikon) for continuous maintenance of focus, all controlled by NIS-Elements software. GFP fluorescence was excited using a 488nm laser line (50mW) and collected with a 525/50 emission filter. Images were acquired with a DU-897 EM-CCD camera (Andor).

Images were brightfield and contrast adjusted using ImageJ (v2.0; National Institutes of Health, Bethesda, MD) and Photoshop CS3 (v10.0, Adobe), and figures were compiled in Illustrator CC (2015.2, Adobe).

### Image and data analysis

For peroxisome and endosome line-scan distribution measurements, maximum-intensity projections of fluorescence micrographs and brightfield images (for hyphae) were obtained using ImageJ. Brightfield images were traced, using the segmented line tool (line width 25), from the hyphal tip and fluorescence micrographs were used to project the average fluorescence intensity. For normalization in Figures 3 and 4, each condition’s average intensity values at each point along hyphae were normalized against its baseline average (10–25 μm). For flux measurements, the number of puncta crossing a line 10 μm perpendicular to and from the hyphal tip was calculated. Endosome, peroxisome, and PxdA velocities and run-lengths were measured using ImageJ. Maximum-intensity projections were generated from time-lapse sequences to define the trajectory of particles of interest. The segmented line tool was used to trace the trajectories and map them onto the original video sequence, which was subsequently resliced to generate a kymograph. The instantaneous velocities of individual particles were calculated from the inverse of the slopes of kymograph traces, and run-length measurements were obtained from each of these traces. For measurements of normalized intensity during colocalized runs between PxdA/endosomes and peroxisomes (Fig. 2 E and Fig. 3 E), peroxisomes and associated puncta (i.e., PxdA or endosomes) were traced using ImageJ at a point along the trajectory of the path (in the direction of movement) that was clear and distinct from other puncta on the micrograph. Line intensity profiles for each channel individually were generated. The “0” along the X-axis corresponds to the position of peak intensity for the peroxisome signal. Intensities for each channel were normalized to that channel’s peak intensity for each co-migrating run.

Data visualization and statistical analyses were performed using GraphPad Prism (6.0d; GraphPad Software), Excel (v14.5.8; Microsoft), and ImageJ (2.0).

### Online Supplemental Material

Figure S1 shows that peroxisomes accumulate in the hyphal tip in PxdA mutant and deletion strains, but early endosomes, nuclei, and cytoplasmic dynein are unaffected. Figure S2 shows that tagging PxdA does not affect its function and that expressing its coiled-coil domain alone moves similarly to the full-length protein. Video 1 shows peroxisome (PTS1-mCherry) dynamics in a wild-type hypha. Video 2 shows peroxisome (PTS1-mCherry) dynamics are perturbed in a *pxdA*Δ hypha. Video 3 shows endosome dynamics in a wild-type hypha. Video 4 shows that endosome dynamics are unaffected in a *pxdA*Δ hypha. Video 5 shows PxdA-GFP dynamics in a hypha. Video 6 shows PxdA-GFP co-migrating with a moving peroxisome. Video 7 shows PxdA-GFP colocalizing with early endosomes. Video 8 shows an early endosome co-migrating with a moving peroxisome. Table S1 lists the *A. nidulans* strains used in this study.

## Acknowledgements

We thank the Reck-Peterson lab and Andres Leschziner for critical feedback on experiments and the manuscript. We thank Jennifer Waters, Talley Lambert, and The Nikon Imaging Center at Harvard Medical School for helpful technical advice and imaging training. S.L.RP is supported by National Institutes of Health grants R01-GM107214 and R01-GM100947

**Abbreviations List**
PxdA,: peroxisome distribution mutant A
CC,: coiled-coil domain
UR,: uncharacterized region
WT,: wild-type
FL,: full-length

## Supplemental Material

### Supplemental Table

**Table S1.**
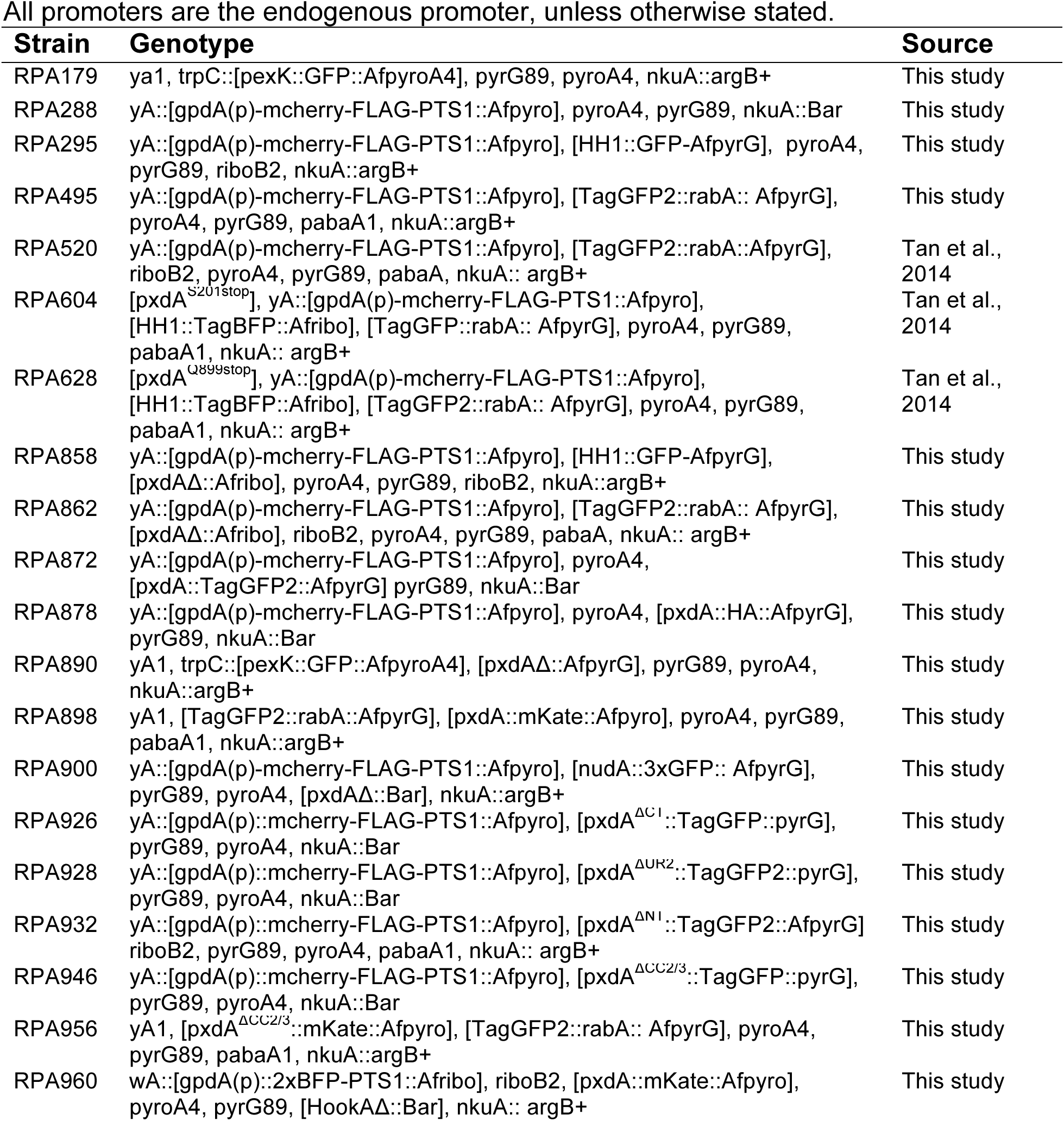
*A. nidulans* strains used in this study

### Supplemental Figure Legends

**Figure S1.**
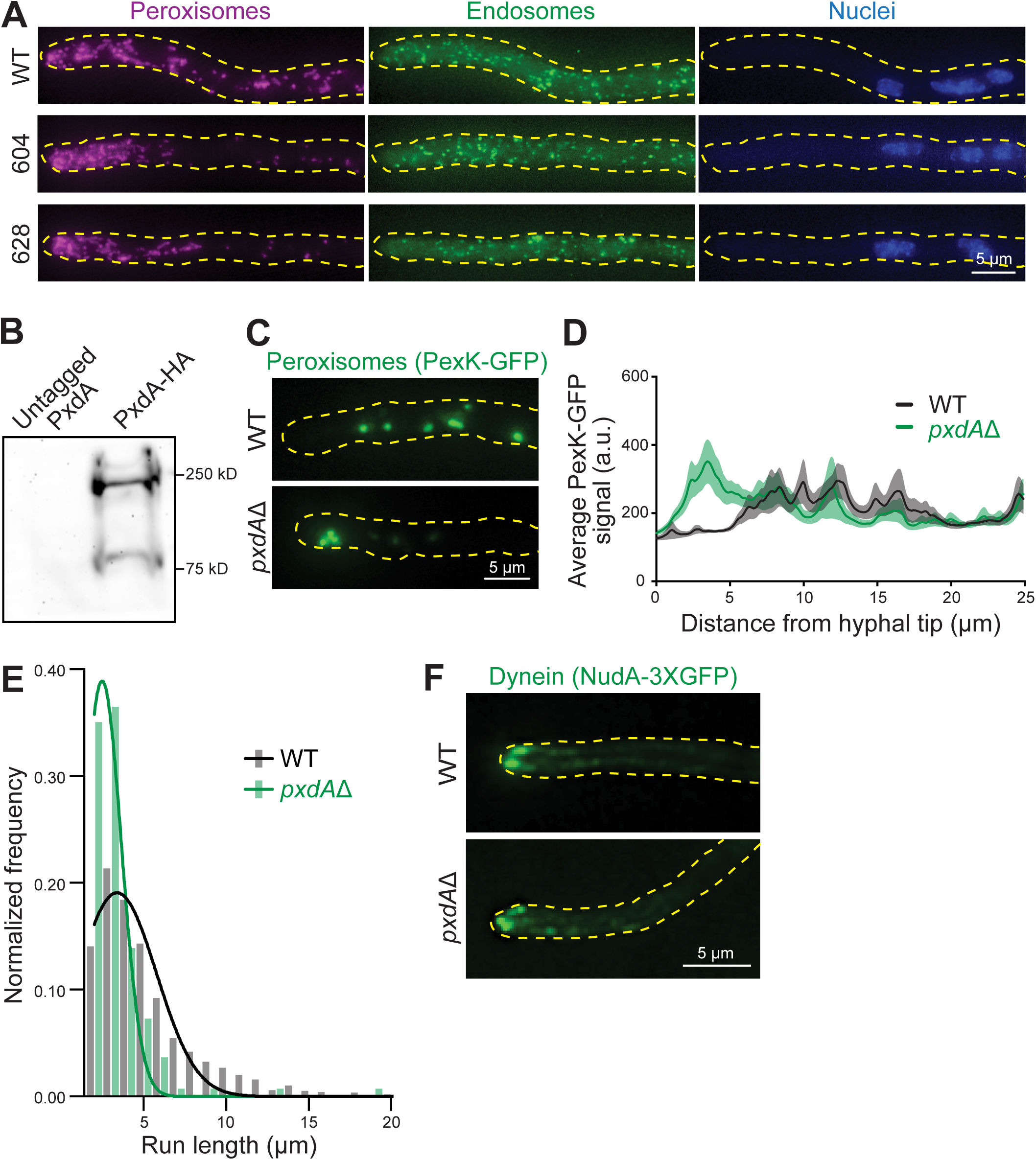
*pxdA* is required for peroxisome movement and distribution, but not for the distribution of dynein. A) Representative micrographs of hyphae labeled with PTS1-mCherry peroxisomes, GFP-RabA/5a endosomes, and HH1-TagBFP nuclei from wild-type (WT; top) and two mutant strains isolated in our original screen (Tan et al., 2014): RPA604 (middle), and RPA628 (bottom). Scale bar = 5 μm. B) Lysates of *A. nidulans* cells expressing PxdA-HA and a non-HA tag control run on SDS-PAGE and immunoblotted with a mouse anti-HA (Sigma) antibody. C) Representative micrographs of maximum-intensity projections of PexK/Pex11-GFP-labeled peroxisomes along hyphae from WT and *pxdA*Δ strains. Scale bar = 5 μm. D) PexK/Pex11-GFP distribution was quantified using line scans of fluorescence micrographs and displayed as the mean (solid lines) +/− SEM (shading) fluorescence intensity as a function of distance from the hyphal tip. Distribution near the hyphal tip was significantly different between genotypes (p < 0.001, two-way ANOVA, Bonferroni post-hoc test significant between 3.14 μm and 4.22 μm from hyphal tip, n = 28 [WT], n = 32 [*pxdA*Δ]). E) Histogram of peroxisome run lengths comparing WT versus *pxdA*Δ were calculated from kymographs generated from 1-min time-lapse movies. The X-axis label begins at 2 μm, the minimum run length we could confidently detect. The distribution of run lengths is statistically different (p < 0.0001, Kolmogorov-Smirnov test, n = 1195 [WT], n = 136 [*pxdA*Δ]). F) Representative micrographs of maximum-intensity projections of nudA-3xGFP-labeled cytoplasmic dynein “comets” along hyphae from WT and *pxdA*Δ strains. Scale bar = 5 μm.

**Figure S2.**
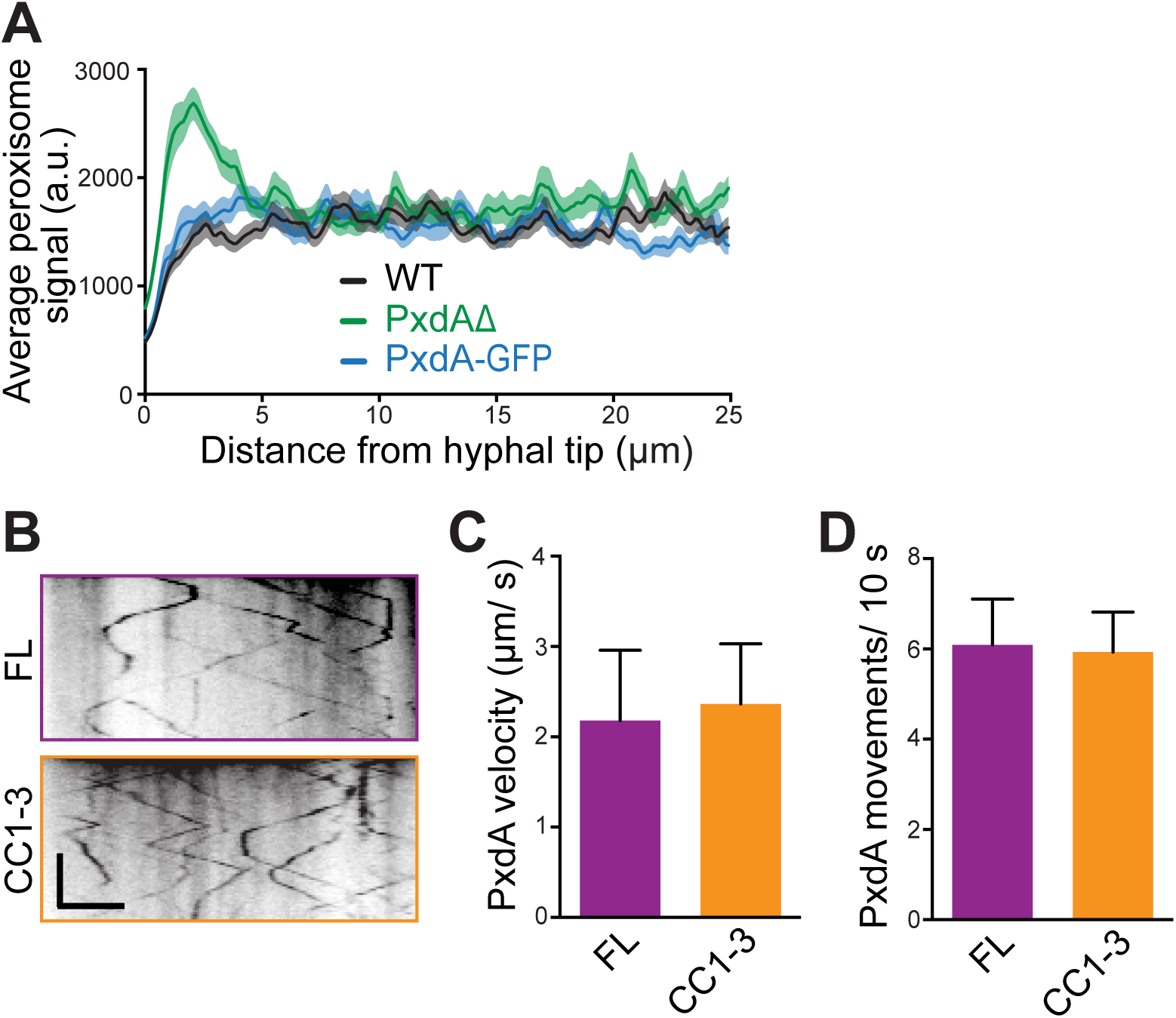
The coiled-coil region of PxdA is necessary and sufficient for movement. A) Line scans of peroxisome (PTS1-mCherry) distribution along hyphae from wild-type (WT), PxdAΔ, and PxdA-GFP strains. B) Representative kymographs of full-length (FL) and CC1-3 PxdA-GFP. C) Bar graph of PxdA-GFP velocities calculated from kymographs of 10 s time-lapse movies. PxdA velocities were 2.18 +/− 0.78 (SD) μm/ s for FL PxdA (n = 151) and 2.37 +/− 0.66 μm/ s for CC1-3 PxdA (n = 188). p < 0.01, Kolmogorov-Smirnov test. D) Bar graph of the number of PxdA-TagGFP movements calculated as the number of puncta crossing a line drawn perpendicular to hyphae during a 10 s time-lapse video. FL PxdA movements were 6.09 +/− 1.01 (SEM; n = 11) and CC1-3 movements were 5.93 +/− 0.89 (n = 14) (p = 0.9049, Student’s t-test).

### Video Legends

**Video 1. Peroxisome movement in a wild-type hypha.** Peroxisome (PTS1-mCherry) movement and dynamics in a wild-type *A. nidulans* hypha. Images were analyzed by time-lapse epifluorescence microscopy using a Deltavision Core microscope (GE Healthcare Life Sciences). Frames were taken every 500 ms for 1 min. Video frame rate is 10 frames/s.

**Video 2. Peroxisome movement is perturbed in *pxdA*Δ hyphae.** Peroxisome (PTS1-mCherry) movement and dynamics in a *pxdA*Δ hypha. Images were analyzed by time-lapse epifluorescence microscopy using a Deltavision Core microscope (GE Healthcare Life Sciences). Frames were taken every 500 ms for 1 min. Video frame rate is 10 frames/s.

**Video 3. Endosome movement in wild-type hyphae.** Endosome (GFP-RabA/5a) movement and dynamics in a wild-type hypha. Images were analyzed by time-lapse epifluorescence microscopy using a Nikon Ti microscope. Frames were taken every 150 ms for 10 s. Video frame rate is 15 frames/ s. Similar videos were used for kymographs generated in Fig. 1 F.

**Video 4. Endosome movement in *pxdA*Δ hyphae.** Endosome (GFP-RabA/5a) movement and dynamics in a *pxdA*Δ hypha. Images were analyzed by time-lapse epifluorescence microscopy using a Nikon Ti microscope. Frames were taken every 150 ms for 10 s. Video frame rate is 15 frames/ s. Similar videos were used for kymographs generated in Fig. 1 F.

**Video 5. PxdA protein exhibits rapid, bidirectional movement.** PxdA-GFP dynamics along a hypha. Images were analyzed by time-lapse epifluorescence microscopy using a Nikon Ti microscope. Frames were taken every 100 ms for 15 s. Video frame rate is 15 frames/ s. This video was used to generate the kymograph in Fig. 2 A.

**Video 6. PxdA comigrates at the leading edge of moving peroxisomes.** PxdA-GFP (green) and PTS1-mCherry-labeled peroxisome (magenta) dynamics in a wild-type hypha. Both channels were acquired simultaneously and analyzed by time-lapse epifluorescence microscopy using a Deltavision OMX microscope (GE Healthcare Life Sciences). Frames were taken every 250 ms for 13.75 s. Video frame rate is 15 frames/ s. This video was used to generate the kymograph in Fig. 2 D (left panel). Scale bar = 5 μm.

**Video 7. PxdA colocalizes with early endosomes.** GFP-RabA/5a-endosomes (green) and PxdA-mKate (magenta) in a wild-type hypha. Both channels were acquired simultaneously and analyzed by time-lapse epifluorescence microscopy using a Deltavision OMX microscope (GE Healthcare Life Sciences). Frames were taken every 250 ms for 7.5 s. Video frame rate is 5 frames/ s.

**Video 8. Early endosomes comigrate with moving peroxisomes.** GFP-RabA/5a-labeled endosomes (green) and PTS1-mCherry-labeled peroxisomes (magenta) in a wild-type hypha. White arrows indicate comigrating puncta. Both channels were acquired simultaneously and analyzed by time-lapse epifluorescence microscopy using a Deltavision OMX microscope (GE Healthcare Life Sciences). Frames were taken every 150 ms for 10.5 s. Video frame rate is 15 frames/ s. Scale bar = 5 μm. This video was used to generate kymograph in Fig. 3 D (left panel).

## References

Abenza, J.F., A. Pantazopoulou, J.M. Rodriguez, A. Galindo, and M.A. Penalva. 2009. Long-distance movement of Aspergillus nidulans early endosomes on microtubule tracks. Traffic. 10: 57–75.

Baumann, S., T. Pohlmann, M. Jungbluth, A. Brachmann, and M. Feldbrugge. 2012. Kinesin-3 and dynein mediate microtubule-dependent co-transport of mRNPs and endosomes. J Cell Sci. 125: 2740–2752.

Bharti, P., W. Schliebs, T. Schievelbusch, A. Neuhaus, C. David, K. Kock, C. Herrmann, H.E. Meyer, S. Wiese, B. Warscheid, C. Theiss, and R. Erdmann. 2011. PEX14 is required for microtubule-based peroxisome motility in human cells. J Cell Sci. 124: 1759–1768.

Bielska, E., M. Schuster, Y. Roger, A. Berepiki, D.M. Soanes, N.J. Talbot, and G. Steinberg. 2014. Hook is an adapter that coordinates kinesin-3 and dynein cargo attachment on early endosomes. J Cell Biol. 204: 989–1007.

Cianfrocco, M.A., M.E. DeSantis, A.E. Leschziner, and S.L. Reck-Peterson. 2015. Mechanism and Regulation of Cytoplasmic Dynein. Annu Rev Cell Dev Biol. 31: 83–108.

Egan, M.J., M.A. McClintock, I.H. Hollyer, H.L. Elliott, and S.L. Reck-Peterson. 2015. Cytoplasmic dynein is required for the spatial organization of protein aggregates in filamentous fungi. Cell Rep. 11: 201–209.

Egan, M.J., M.A. McClintock, and S.L. Reck-Peterson. 2012a. Microtubule-based transport in filamentous fungi. Curr Opin Microbiol. 15:637–645. 23

Egan, M.J., K. Tan, and S.L. Reck-Peterson. 2012b. Lis1 is an initiation factor for dynein-driven organelle transport. J Cell Biol. 197: 971–982.

Fu, M.M., and E.L. Holzbaur. 2014. Integrated regulation of motor-driven organelle transport by scaffolding proteins. Trends Cell Biol.

Gibson, D.G., L. Young, R.Y. Chuang, J.C. Venter, C.A. Hutchison, 3rd, and H.O. Smith. 2009. Enzymatic assembly of DNA molecules up to several hundred kilobases. Nat Methods. 6: 343–345.

Glater, E.E., L.J. Megeath, R.S. Stowers, and T.L. Schwarz. 2006. Axonal transport of mitochondria requires milton to recruit kinesin heavy chain and is light chain independent. J Cell Biol. 173: 545–557.

Gohre, V., E. Vollmeister, M. Bolker, and M. Feldbrugge. 2012. Microtubule-dependent membrane dynamics in Ustilago maydis: Trafficking and function of Rab5a-positive endosomes. Commun Integr Biol. 5: 485–490.

Griffith, J., M.A. Penalva, and F. Reggiori. 2011. Adaptation of the Tokuyasu method for the ultrastructural study and immunogold labelling of filamentous fungi. Journal of Electron Microscopy. 60: 211–216.

Guimaraes, S.C., M. Schuster, E. Bielska, G. Dagdas, S. Kilaru, B.R. Meadows, M. Schrader, and G. Steinberg. 2015. Peroxisomes, lipid droplets, and endoplasmic reticulum “hitchhike” on motile early endosomes. J Cell Biol.

Harada, A., Y. Takei, Y. Kanai, Y. Tanaka, S. Nonaka, and N. Hirokawa. 1998. Golgi vesiculation and lysosome dispersion in cells lacking cytoplasmic dynein. J Cell Biol. 141:51–59. 24

Higuchi, Y., P. Ashwin, Y. Roger, and G. Steinberg. 2014. Early endosome motility spatially organizes polysome distribution. J Cell Biol. 204: 343–357.

Horio, T., and B. Oakley. 2005. The role of microtubules in rapid hyphal tip growth of Aspergillus nidulans. Mol Biol Cell. 16: 918–926.

Hynes, M.J., S.L. Murray, G.S. Khew, and M.A. Davis. 2008. Genetic analysis of the role of peroxisomes in the utilization of acetate and fatty acids in Aspergillus nidulans. Genetics. 178: 1355–1369.

Kardon, J.R., and R.D. Vale. 2009. Regulators of the cytoplasmic dynein motor. Nat Rev Mol Cell Biol. 10: 854–865.

Kural, C., H. Kim, S. Syed, G. Goshima, V.I. Gelfand, and P.R. Selvin. 2005. Kinesin and dynein move a peroxisome in vivo: a tug-of-war or coordinated movement? Science. 308: 1469–1472.

Lee, S.B., and J.W. Taylor. 1990. Isolation of DNA from fungal mycelia and single spores. *In* PCR Protocols: A Guide to Methods and Applications D.G. M. Innis, J. Sninsky and T. White, eds., editor. Academic Press, Orlando, Florida. 282–287.

Liu, F., S.K. Ng, Y. Lu, W. Low, J. Lai, and G. Jedd. 2008. Making two organelles from one: Woronin body biogenesis by peroxisomal protein sorting. J Cell Biol. 180: 325–339.

Maday, S., and E.L. Holzbaur. 2012. Autophagosome assembly and cargo capture in the distal axon. Autophagy. 8: 858–860.

McKenney, R.J., W. Huynh, M.E. Tanenbaum, G. Bhabha, and R.D. Vale. 2014. Activation of cytoplasmic dynein motility by dynactin-cargo adapter complexes. Science. 345: 337–341.

Murk, J.L., G. Posthuma, A.J. Koster, H.J. Geuze, A.J. Verkleij, M.J. Kleijmeer, and B.M. Humbel. 2003. Influence of aldehyde fixation on the morphology of endosomes and lysosomes: quantitative analysis and electron tomography. J Microsc. 212: 81–90.

Nayak, T., E. Szewczyk, C.E. Oakley, A. Osmani, L. Ukil, S.L. Murray, M.J. Hynes, S. Osmani, and B. Oakley. 2006. A versatile and efficient genetargeting system for Aspergillus nidulans. Genetics. 172: 1557–1566.

Neuhaus, A., C. Eggeling, R. Erdmann, and W. Schliebs. 2015. Why do peroxisomes associate with the cytoskeleton? Biochim Biophys Acta.

Rapp, S., R. Saffrich, M. Anton, U. Jakle, W. Ansorge, K. Gorgas, and W.W. Just. 1996. Microtubule-based peroxisome movement. J Cell Sci. 109 (Pt 4):837–849.

Roghi, C., and V.J. Allan. 1999. Dynamic association of cytoplasmic dynein heavy chain 1a with the Golgi apparatus and intermediate compartment. J Cell Sci. 112 (Pt 24):4673–4685.

Schrader, M., S.J. King, T.A. Stroh, and T.A. Schroer. 2000. Real time imaging reveals a peroxisomal reticulum in living cells. J Cell Sci. 113 (Pt 20):3663–3671.

Shcherbo, D., C.S. Murphy, G.V. Ermakova, E.A. Solovieva, T.V. Chepurnykh, A.S. Shcheglov, V.V. Verkhusha, V.Z. Pletnev, K.L. Hazelwood, P.M. Roche, S. Lukyanov, A.G. Zaraisky, M.W. Davidson, and D.M. Chudakov. 2009. Far-red fluorescent tags for protein imaging in living tissues. Biochem J. 418: 567–574.

Straubinger, B., Straubinger, E., Wirsel, S., Turgeon, G., and Yoder, O.C. 1992. Versitile fungal transformation vectors carrying the selectable bar gene of Streptomyces hygroscopicus. Fungal Genetics Newsletter. 39: 82–83.

Subach, O.M., I.S. Gundorov, M. Yoshimura, F.V. Subach, J. Zhang, D. Gruenwald, E.A. Souslova, D.M. Chudakov, and V.V. Verkhusha. 2008. Conversion of red fluorescent protein into a bright blue probe. Chem Biol. 15: 1116–1124.

Szewczyk, E., T. Nayak, C.E. Oakley, H. Edgerton, Y. Xiong, N. Taheri-Talesh, S. Osmani, B. Oakley, and B. Oakley. 2006. Fusion PCR and gene targeting in Aspergillus nidulans. Nat Protoc. 1: 3111–3120.

Tan, K., A.J. Roberts, M. Chonofsky, M.J. Egan, and S.L. Reck-Peterson. 2014. A microscopy-based screen employing multiplex genome sequencing identifies cargo-specific requirements for dynein velocity. Mol Biol Cell. 25: 669–678.

Tanaka, Y., Y. Kanai, Y. Okada, S. Nonaka, S. Takeda, A. Harada, and N. Hirokawa. 1998. Targeted disruption of mouse conventional kinesin heavy chain, kif5B, results in abnormal perinuclear clustering of mitochondria. Cell. 93: 1147–1158.

Vale, R.D. 2003. The molecular motor toolbox for intracellular transport. Cell. 112:467–480. 27

van Spronsen, M., M. Mikhaylova, J. Lipka, M.A. Schlager, D.J. van den Heuvel, M. Kuijpers, P.S. Wulf, N. Keijzer, J. Demmers, L.C. Kapitein, D. Jaarsma, H.C. Gerritsen, A. Akhmanova, and C.C. Hoogenraad. 2013. TRAK/Milton motor-adaptor proteins steer mitochondrial trafficking to axons and dendrites. Neuron. 77: 485–502.

Wang, X., and T.L. Schwarz. 2009. The mechanism of Ca2+ -dependent regulation of kinesin-mediated mitochondrial motility. Cell. 136: 163–174.

Wedlich-Soldner, R., A. Straube, M.W. Friedrich, and G. Steinberg. 2002. A balance of KIF1A-like kinesin and dynein organizes early endosomes in the fungus Ustilago maydis. EMBO J. 21: 2946–2957.

Xiang, X., S.M. Beckwith, and N.R. Morris. 1994. Cytoplasmic dynein is involved in nuclear migration in Aspergillus nidulans. Proc Natl Acad Sci U S A. 91: 2100–2104.

Xiang, X., C. Roghi, and N.R. Morris. 1995. Characterization and localization of the cytoplasmic dynein heavy chain in Aspergillus nidulans. Proc Natl Acad Sci U S A. 92: 9890–9894.

Zhang, J., R. Qiu, H.N. Arst, Jr., M.A. Penalva, and X. Xiang. 2014. HookA is a novel dynein-early endosome linker critical for cargo movement in vivo. J Cell Biol. 204: 1009–1026.

Zhang, J., L. Zhuang, Y. Lee, J.F. Abenza, M.A. Penalva, and X. Xiang. 2010. The microtubule plus-end localization of Aspergillus dynein is important for dynein-early-endosome interaction but not for dynein ATPase activation. J Cell Sci. 123: 3596–3604.

